# The Role of Meningeal Lymphatic Vessels and Perivascular Cerebrospinal Fluid Flow in Age-Related Processing Speed Decline

**DOI:** 10.64898/2026.04.05.716121

**Authors:** Micaela Andreo, Dinesh K Sivakolundu, Mark Zuppichini, Kathryn West, Jeffrey Spence, Susan A Gauthier, Thanh Nguyen, Bart Rypma

## Abstract

Meningeal lymphatic vessels (mLV) play essential roles in draining cerebrospinal fluid (CSF) into peripheral blood. The mLVs are hypothesized to be supportive structures to the glymphatic system, which is thought to remove metabolic byproducts from brain parenchyma and has been most directly studied in rodent models. Previous rodent studies have indicated a correlation between mLV function and cognitive performance, but this relationship in humans remains unexplored. Age-related declines in glymphatic system efficiency in humans and cognitive performance have been observed separately. This study investigates age- and sex-related differences in CSF production via choroid plexus volumes, mLV characteristics, and glymphatic system efficiency, overall elucidating the implication of cerebral lymphatic function on cognition. We recruited 26 healthy adults from Dallas-Fort Worth and acquired magnetic resonance images. mLVs along the sagittal sinus were visualized and segmented from T_2_-FLAIR images. The glymphatic system was evaluated by measuring diffusivity along the perivascular space. Choroid plexus volume and brain volume were estimated from T_1_-MPRAGE. Neuropsychological tests were conducted to assess cognitive function. Our findings indicate that glymphatic function diminishes with age, while mLV and choroid plexus volumes increase. Males displayed greater mLV volume than females, yet no sex differences were found in glymphatic function or choroid plexus volume. Notably, mLV volume increased as glymphatic function declined, independent of age. Moreover, a glymphatic-mLV latent variable significantly predicted processing speed, underscoring the influence of cerebral lymphatics on cognition. In conclusion, this study highlights a decline in glymphatic function with age, accompanied by increased mLV volumes and altered processing speed. These lymphatic system changes may underlie or contribute to the cognitive declines observed in healthy and pathological aging.

**Significance Statement:** The glymphatic system and meningeal lymphatic vessels play crucial roles in removing brain cell waste. The relationship between these systems and their effect on human cognition, particularly processing speed, is unknown. We demonstrate that these systems change with advancing age. Variations in cerebral lymphatic function contribute to differences in processing speed independent of age, ultimately affecting higher-order cognitive function. The findings presented have implications for cognitive function in both healthy and diseased states.

## Introduction

The glymphatic system and meningeal lymphatic vessels (mLV) play essential roles in clearing cellular waste from the brain and draining cerebrospinal fluid (CSF) into peripheral blood. The glymphatic system removes cellular byproducts from the brain interstitium and transfers them into the CSF (Iliff et al., 2012), while the mLVs transfer these byproducts from the CSF into the peripheral blood (Aspelund et al., 2015; Louveau et al., 2015). While some work has been done, precise relationships between the glymphatic system processes and cognition in the human model are not yet completely understood. There are several theories of glymphatic system physiology and aging (Nedergaard & Goldman, 2020; Verheggen et al., 2018). One such theory proposes that age-related disruption of the CSF flow through brain parenchyma, possibly related to sleep dysfunction, leads to subsequent metabolic waste accumulation and neural dysfunction (Dagum et al., 2026; Alshammari et al., 2026; Hein et al., 2026).

In the intact system, CSF produced via the choroid plexus circulates in the central nervous system (CNS) through the ventricles and spinal cord, eventually entering the subarachnoid space (Jessen et al., 2015). CSF from the subarachnoid space enters the brain parenchyma via peripenetrating arteriolar space, allowing for exchange with brain interstitial fluid that is mediated by glial cells (Iliff et al., 2012). The mixed fluid-carrying waste then exits the perivascular space and returns to the subarachnoid space, forming the glymphatic system (Iliff et al., 2012). CSF is then reabsorbed into the bloodstream through various routes, including the mLVs (Aspelund et al., 2015; Louveau et al., 2015). It is unclear how age-related differences in cerebral lymphatics affect brain function and cognition.

Previous research in both rodents and humans have implicated glymphatic system changes in the pathophysiology of healthy aging (Jessen et al., 2015), various neurologic disorders such as Alzheimer’s disease (Kress et al., 2014; Taoka et al., 2017), traumatic brain injury (Jessen et al., 2015), stroke (Mestre et al., 2020), small vessel disease (Zhang et al., 2021), and multiple sclerosis (Carotenuto et al., 2022), as well as diabetes (Jiang et al., 2017) and obstructive sleep apnea (Lee et al., 2022). In Alzheimer’s disease and small vessel disease, suppressed glymphatic transport contributes to cognitive impairment and dementia (Wang et al., 2023; Da Mesquita et al., 2018; Zhang et al., 2021). The mechanisms through which glymphatic alterations lead to cognitive impairments in humans remain unclear. A number of mechanisms by which glymphatic alterations could lead to cognitive impairment have been proposed, including pathological protein deposition (amyloid and tau) that increases inflammation and neurotoxicity, as well as iron deposition that could disrupt neural function (Zhou et al., 2020; Iliff et al., 2012; Carare et al., 2020; Li et al., 2022). Such disruptions might exert effects on neural transmission efficiency, spike train frequency, and cognitive processing speed (Zhao et al., 2021). Rodent studies suggest that impaired mLV function is related to cognitive impairment in aging (Ahn et al., 2019; Da Mesquita et al., 2018) Alzheimer’s disease (Wang et al., 2019), and Parkinson’s disease (Ding et al., 2021). Restoring mLV function improved cognition in an Alzheimer’s disease model by improving CSF drainage (Ding et al., 2021). To our knowledge, no studies have investigated the role of mLVs in basic aspects of human cognition, such as processing speed and working memory. Thus, we hypothesize that differences in cerebral lymphatic function contribute to the processing speed variability seen in healthy humans, which in turn affects higher-order cognitive functioning.

## 2. Materials and Methods

### 2.1 Ethics Statement

The study was approved by the University of Texas Southwestern Medical Center (UTSW) Institutional Review Board. Informed written consent was obtained from all participants prior to study participation.

### 2.2 Research Participants

The study group was comprised of 26 healthy individuals recruited using advertisements and flyers distributed throughout the Dallas-Fort Worth Metroplex area. Inclusion criteria for all participants were (i) male or female participants between the ages of 18 and 65. Exclusion criteria for all participants included (i) left-handed participants, (ii) pregnant or nursing women, (iii) history of smoking or cardiopulmonary illness, (iv) prior history of neuropsychiatric conditions, and (v) contraindications to MRI scanning.

### 2.3 MRI Data Acquisition

MRI scans were performed on a 3T MRI scanner (Philips Medical Systems, Best, The Netherlands) equipped with a 32-channel phased array head coil at the UTSW Advanced Imaging Research Center. Participants underwent high-resolution T_2_-weighted fluid attenuated inversion recovery (T_2_ FLAIR), T_1_-weighted magnetization-prepared rapid acquisition gradient-echo (MPRAGE) images, susceptibility-weighted imaging (SWI), and diffusion tensor imaging (DTI).

MPRAGE was acquired with the following scan parameters: Repetition time (TR)=8.1ms, echo time (TE)=3.7ms, shot interval=2100ms, inversion time (TI)=1100ms, resolution=1mm^3^ isotropic, flip angle =12°, field of view (FOV)=256mm × 204mm × 160mm, matrix size=256 × 204 × 160, sagittal slice orientation.

T_2_-FLAIR was acquired with the following scan parameters: TR=4800ms TE=344ms, TI=1600ms, FOV=250mm × 250mm × 179 mm, matrix size=228 × 227 × 163, resolution=1.1mm^3^ isotropic, sagittal slice orientation.

SWI was acquired with the following scan parameters: field of view = 230mm × 180mm x 140mm, no of echoes = 4, echo times = 6:12:18:24, voxel size = 0.6 x 1, flip angle = 15° both magnitude and phase images for each echo were saved.

DTI was acquired with the following scan parameters: Single-shot echo planar imaging (EPI) sequence with TR=6500ms, TE=62ms, FOV=224 x 224mm^2^, resolution=2.0 x 2.0mm^2^, slice thickness=2.20mm, number of slices=62, slice gap=0mm, SENSE-reduction factor=2.3, with b-shells of b = 0 s/mm^2^, 1000s/mm^2^ across 30 directions.

### 2.4 Meningeal Lymphatic Vessel (mLV) visualization, segmentation, and phenotyping

#### 2.4.1. mLV visualization

In humans, mLV are typically observed along the dural venous sinuses in the dorsal regions and along the cranial nerves in the ventral regions (Absinta et al., 2017). Three-dimensional FLAIR sequence permits visualization of the mLV based on the internal signal from protein-rich lymphatic fluid without the need for any external contrast agents (Albayram et al., 2022). Protein concentration in lymph is usually in the range of ∼2-4g/dl that includes Albumin (concentration in lymph is 1.6-1.8g/dl) and other proteinaceous components related to cells, solutes, and debris. A phantom study examining various FLAIR scan parameters found that mLVs corresponding to this protein concentration can be effectively visualized when the parameters were set at TR=5000ms, TE=386ms, and TI=1400ms (Albayram et al., 2022). We used similar scan parameters in our study to visualize mLVs (see above for scan parameters, see Fig 1B).

**Fig 1.**
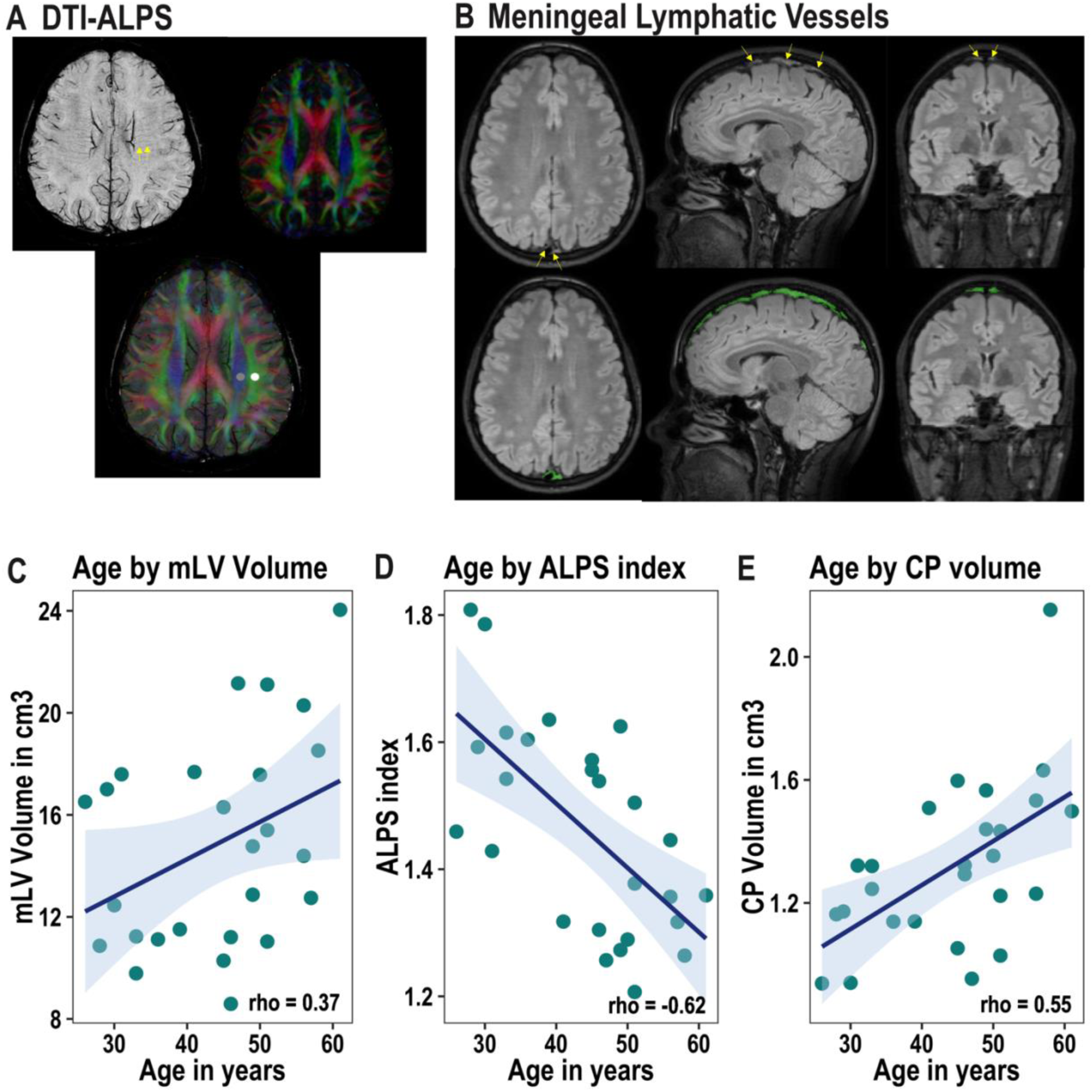
A. Diffusion Tensor Imaging along the Perivascular Space (DTI-ALPS) for Glymphatic System Efficiency Assessment. The left image is a minimal intensity projection of the Susceptibility-Weighted Imaging, showing medullary veins perpendicular to the lateral ventricular wall (indicated by arrows). The image on the right is a diffusion tensor image, with blue fibers representing projection fibers running in the superior-inferior direction and green fibers representing superior longitudinal fibers (association fibers) running in the anterior-posterior direction. The overlaid image below demonstrates the perpendicularity of the medullary veins and the major fibers, with spheres indicating the region of interest used for calculating the ALPS index. B. mLV along the Dural Venous Sinus. The image shows the mLV along the dural venous sinus, with arrows and a shaded region in green indicating the location. C. Scatter plot depicting the relationship between mLV volume and Age. D. Scatter plot depicting the relationship between ALPS index and Age. E. Scatter plot depicting the relationship between choroid plexus volume and Age.

Using these scan parameters, Albayram and colleagues validated the technique in humans and found that mLVs along the anterior and middle portions of sagittal sinus can be observed across all participants with inter-rater reliability of 1 (Albayram et al., 2022). However, mLVs along other parts of the dural venous sinuses were visible in only 58.0% - 99.7% of participants, depending on the specific location and positioning of the participant within the scanner. Due to inconsistency in visualizing mLVs across the various dural venous sinuses, excluding the sagittal sinus, we decided to focus solely on characterizing the mLVs along the sagittal sinus in our study. In addition, ventral mLVs were excluded in our study as our voxel size of 1.1mm^3^ was too large to consistently visualize the ventral mLVs.

#### 2.4.2. mLV segmentation and phenotyping

mLVs along the sagittal sinus were identified in the FLAIR images for each participant. To assist in the segmentation of mLVs, a template was generated from individual FLAIR images using the ANTs multivariate template construction algorithm (Avants et al., 2011). In the FLAIR template, mLVs were identified and manually segmented. During the segmentation process, there was unavoidable contamination from blood and CSF in the venous sinus and subarachnoid space. This contamination was addressed by removing voxels within the mask with intensities lower than the 90th percentile of the CSF mask intensity. To ensure accuracy, the final segmented mLVs from the template were reviewed by three experts in the field (TN, DKS, SAG) to confirm that only the mLVs along the sagittal sinus were included in the segmentation. An inverse transformation was then applied to the template mask, allowing for the conversion of the mLV mask in template space into the native space of each participant. This transformation produced individual mLV masks in native space. Finally, the mLV masks in native space were visually inspected, and any registration errors or inaccuracies were corrected as necessary. The characteristics of the mLV were assessed by their volume, surface area, and thickness. The volume and surface area were estimated using an in-house Python script. mLV thickness was calculated as the average Euclidean distance across the entire mask.

### 2.5 Glymphatic system evaluation using Diffusion Tensor Imaging Along the Perivascular Space (DTI-ALPS)

The integrity of the glymphatic system was assessed using DTI-ALPS (Taoka et al., 2017). The glymphatic system consists of CSF that clears cellular waste products along the perivascular space. In the region of the lateral ventricular body, the medullary veins and surrounding perivascular space run perpendicular to the ventricular wall in the right-left direction (x-axis). Adjacent to the lateral ventricle in the same plane, projection fibers run in the superior-inferior direction (z-axis), while superior longitudinal fibers (association fibers) run in the anterior-posterior direction (y-axis). This perpendicular arrangement of the major fibers allows for the calculation of diffusivity along the perivascular space. This diffusivity can be normalized by comparing it to the perpendicular axes of the major fibers (i.e., y-axis for projection fibers and z-axis for association fibers). The resulting index is known as the ALPS index (equation 1).

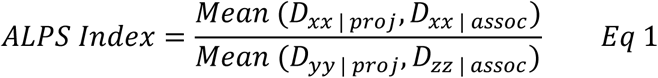

The ALPS index permits in-vivo evaluation of the glymphatic system (Taoka et al., 2017) and has been validated in humans (Zhang et al., 2021). It is reproducible (Tatekawa et al., 2023), and its values correlate with the glymphatic clearance function, which is typically assessed using MRI with the administration of gadolinium-based contrast agents via intrathecal injection (Zhang et al., 2021). A higher ALPS index value indicates greater diffusivity along the perivascular space and more efficient clearance through the glymphatic system. Conversely, a lower ALPS index value indicates less diffusivity along the perivascular space and reduced clearance through the glymphatic system.

To calculate the ALPS index, the following steps were taken. Firstly, the SWI images were processed using the CLEAR-SWI toolkit (Eckstein et al., 2021) and used to identify the medullary vein. Next, diffusion images were corrected for eddy-current distortions and motion using the EDDY tool in the FMRIB Software Library (FSL v5.0.9; Andersson and Sotiropoulos, 2016; Andersson et al., 2016). To account for potential variations in ALPS due to scanner positioning, all participants were checked to ensure they had a neutral head position (Tatekawa et al., 2023). Diffusion tensors were then estimated using the DTIFIT tool. The diffusion images were registered to the magnitude of the first echo of the SWI sequence through rigid body transformation using the FLIRT function in FMRIB’s Linear Image Registration Tool. The resulting registration matrix was applied to perform vector-based registration of the principal diffusion direction field images of each participant to the SWI images using the vecreg function. All transformations were visually inspected to verify proper alignment.

Using the minimal intensity projection map of the SWI images as a reference, along with color-coded principal diffusion direction maps, at least three contiguous slices were selected where the medullary veins ran perpendicular to the association and projection fibers. In the middle slice of the selected three, two three-dimensional spherical regions of interest (ROIs) with a diameter of 3mm were drawn in the right hemisphere: one over the association fibers and the other over the projection fibers (Fig 1A). While the DTI-ALPS was originally calculated using 5-mm spherical regions of interest (ROIs)(Taoka et al., 2017), we chose to draw 3mm to minimize the risk of losing the perpendicularity between fibers and the perivascular space. The ROIs were then transformed back to the diffusion space from the SWI space using the inverse of the initially computed transformation matrix. The transformed ROIs were visually inspected to ensure their correct association with the association and projection fibers. Mean diffusivities in the xx-, yy-, and zz-directions were calculated within the ROIs. Finally, the ALPS index was computed for each participant using Equation 1.

### 2.6 Choroid plexus volume and brain volume estimation

Choroid plexus, cortical grey matter and white matter volumes were estimated from MPRAGE using Freesurfer. The segmented choroid plexus from Freesurfer was visually inspected, and any errors in the segmentation were manually corrected before calculating its volume.

### 2.7 Neuropsychological tests

All participants underwent a comprehensive neuropsychological evaluation with a comprehensive test battery that was used to assess the following cognitive domains: (1) processing speed, measured with Digit-Symbol-Substitution Test and Symbol-Digit-Modalities Test, (2) working memory, measured with backwards digit-span, (3) visuospatial memory, measured with 10/36 spatial-recall test, (4) verbal learning, measured with Selective-Reminding Test, (5) verbal fluency, measured with Controlled-Oral-Word-Association Test, (6) executive function, measured as the difference between the second and first trials in trail making task. The scores from each of these tests were *z*-standardized such that lower z-scores indicated poorer performance, and higher z-scores indicated better performance. The *z*-score for each of the two processing speed tests was averaged to obtain a processing speed composite for every participant.

### 2.8 Statistics

All analyses were conducted using R (version 3.5.2; R Core Team, 2018). To explore associations between continuous variables (such as Age, mLV volume, surface area, thickness, normalized intensity, ALPS index, and neuropsychological test scores), Spearman correlation tests were performed. To examine potential group differences in mLV features between males and females, Wilcoxon rank sum tests were employed. Multivariate regressions were carried out using the Lavaan package (Rosseel, 2012) to investigate relationships among Age, mLV volume, and ALPS index. Additionally, covariance-based structural equation modelling (SEM) was utilized to create a “Lymphatic Latent Variable” from mLV volume and ALPS index, and to explore the relationship between the Lymphatic Latent Variable and neuropsychological test scores. The final models were bootstrapped with 2,000 retrials to aid in interpretation of key path coefficients (recommendation at least 500 or 1,000; see Cheung & Lau, 2009).

## 3. Results

### 3.1 mLV and choroid plexus volume show an increase with age while glymphatic function decreases

As a first aim, we characterized the phenotypic features of the mLVs along the sagittal sinus (Fig 1B) in 26 healthy individuals (range: 26-61 years, mean=44 years, standard deviation=10.4 years, Table 1). We then explored how choroid plexus volume and cerebral lymphatic drainage, assessed via mLV volume and the ALPS index (an indicator of glymphatic function), change with age and sex. mLV volume, r_s_(24)=0.37, p=0.064, and choroid plexus volume, r_s_(24)=0.55, p=0.003, increased with age (Fig 1C, 1E). In contrast, the ALPS index decreased with age, r_s_(24)=-0.62, p<0.001(Fig 1D).

**Table 1:** Phenotypic characteristics of mLV.

| **mLV phenotypic characteristics** | **Mean (Standard deviation)** |
| --- | --- |
| **Volume in cm<sup>3</sup>** | 14.85 (4.10) |
| **Surface area in cm<sup>2</sup>** | 383.40 (75.84) |
| **Average Thickness in mm** | 2.19(0.06) |

### 3.2 Sex differences in mLV volume, ALPS index, and choroid plexus volume

Males had higher mLV volumes compared to females (M_male_=18.1cm^3^, M_female_=12.7cm^3^, W=111, p=0.032) after accounting for differences in intracranial volumes. There were no R-based sex-differences in the ALPS index (p=0.115) and choroid plexus volume (p=0.238).

### 3.3 mLV volume increases as glymphatic function decreases independent of age

Further, we assessed the relationship between mLV and glymphatic function independent of age. Although mLV volume increased with age, this association was not significant after controlling for age-related effects on the ALPS index (ß=0.10, p=0.546, Fig 2A). Notably, the mLV volume increased significantly with decreasing ALPS index (ß=-0.44, p=0.034, Fig 2A), independent of age. There were no significant relationships observed between choroid plexus volume and mLV volume, r_s_(24)=0.05, p=0.807, or choroid plexus volume and ALPS index, r_s_(24)=-0.32, p=0.110.

**Fig 2.**
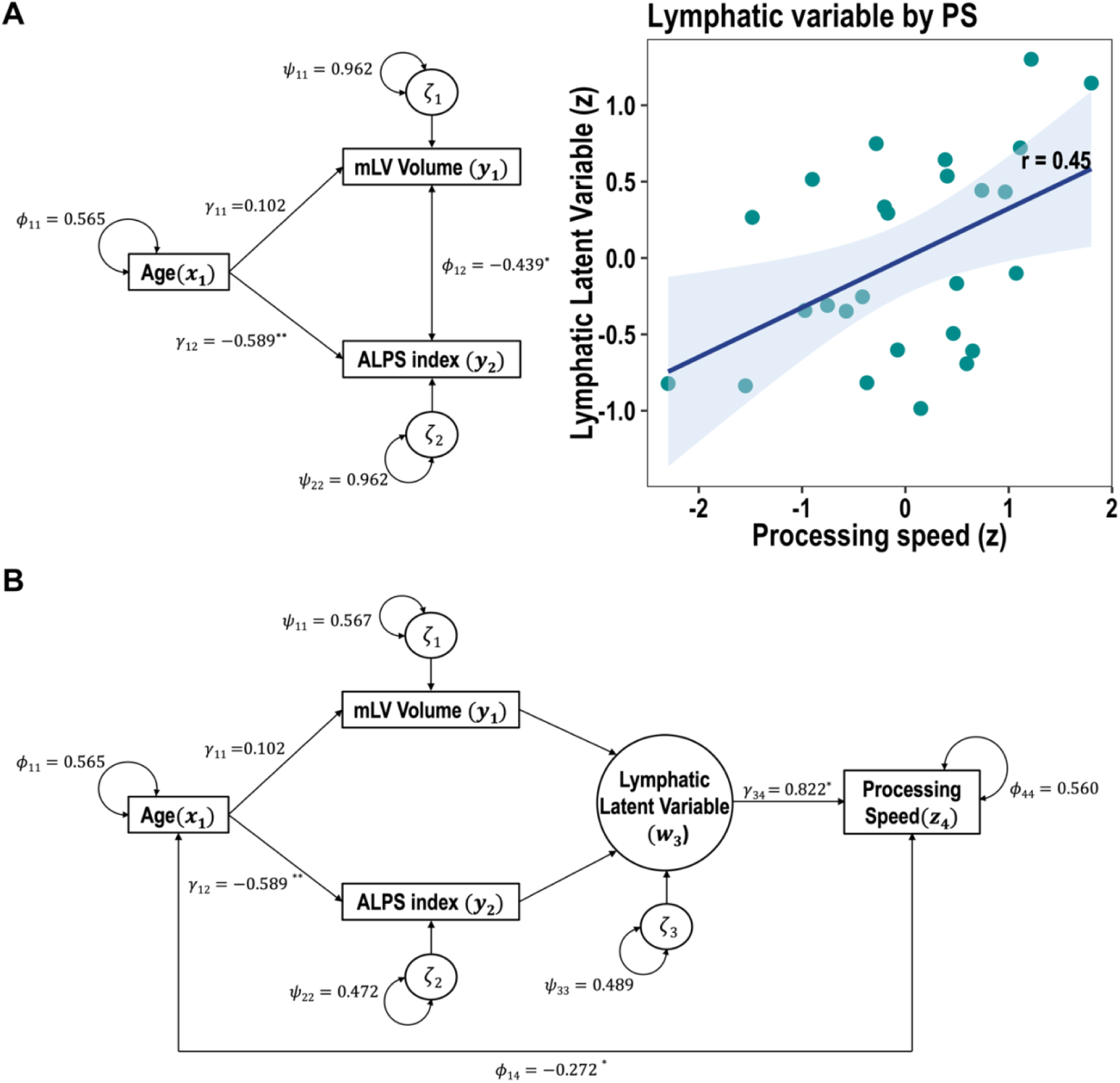
A. Structural equation model used to assess the relationship between Age, mLV volume, and ALPS index. B. Scatter plot depicting the relationship between lymphatic latent variable and processing speed scores. C. Structural equation model used to assess the relationship between Age, Lymphatic Latent Variable, and Processing Speed.

We further investigated the associations between mLV volume, choroid plexus volume, ALPS index, and cortical volume. However, we did not observe any significant relationships between lymphatic and glymphatic variables (mLV volume, choroid plexus volume, ALPS index) and cortical volume (ps>0.05).

### 3.4 Lymphatic latent variable predicts processing speed independent of age and has implications for higher-order cognition

Finally, we investigated how cerebral lymphatic function may impact processing speed, a fundamental cognitive ability known to influence overall cognitive performance. As a proxy for cerebral lymphatic function, we created a latent variable representing the shared variance between mLV volume and the ALPS index (i.e., the lymphatic latent variable; LLV) using SEM. We then looked for an association between the LLV and processing speed independent of age. LLV significantly predicted processing speed (ß=0.82, p=0.045, Fig 2B, and 2C), even after accounting for age-related changes. However, after accounting for LLV, an association between age and processing speed persisted (ß=-0.27, p<0.04, Fig 2C), indicating that factors beyond cerebral lymphatics influence age-related processing speed decline. We did not observe any significant relationships between LLV and other cognitive domains (p>0.05).

The associations we have observed between glymphatic function and processing speed could implicate a critical role for glymphatic waste clearance mechanisms in age-related processing speed decline. Processing speed declines are known to have an impact on higher-order cognitive functions (Salthouse, 1996, 2000; Rypma et al., 2006; Sivakolundu et al., 2019). In fact, in our study, we observed that differences in processing speed explained 32.8% of the variability in visuospatial memory (p=0.002) and 21.8% of the variability in executive function (p=0.017). These findings suggest that alterations in processing speed could influence cognitive domains such as visuospatial memory and executive function.

## 4. Discussion

In this study, we sought to understand how relationships between the mLVs and glymphatic system (DTI-ALPS index and choroid plexus volume) may influence brain function related to cognition, specifically processing speed. Our results showed that the ALPS index, reflecting perivascular CSF flow, decreased with age. These results support the hypothesis that glymphatic function decreases with age (Xiong et al., 2024; Kress et al., 2014). We also found that mLV volume increases with age, suggesting that increases in mLV volume may occur as a compensatory response to age-related decreases in glymphatic function (Jessen et al., 2015; Antila et al., 2017).

Our study aligns with previous scientific literature, both in humans and animals, showing a decline in glymphatic system function and an increase in meningeal lymphatic volume with age (Jessen et al., 2015; Antila et al., 2017). Previous research has demonstrated that mLVs undergo additional growth with age, mediated by a signaling pathway involving vascular endothelial growth factor (VEGF; Antila et al., 2017). It might be that, in response to reduced effectiveness of glymphatic clearance, VEGF secretion is increased in vascular cells, contributing to the enlargement of the mLV volume seen in our sample, along with decreases in age-related glymphatic function.

Although the CSF flow dynamics could affect the cerebral lymphatic system, our findings did not reveal age differences in choroid plexus volume. This result indicates that the volume of choroid plexus, which produces CSF, may not necessarily indicate the extent of CSF production (Jessen et al., 2015). It is also possible that our older sample, consisting of relatively healthy individuals, may be in early stages of the aging process in which lateral ventricles have not enlarged, limiting age-related choroid plexus hypertrophy or increased CSF production.

Our results, showing associations between the lymphatic system (mLV volume and ALPS index) and processing speed performance, support the hypothesis that cerebral lymphatic system function and integrity play important supportive roles in human cognition. Glymphatic clearance declines could lead to cognitive declines, some of which include increased pathological protein and iron depositions, which could lead to subsequent inflammation and neural dysfunction (Zhou et al., 2020; Iliff et al., 2012; Carare et al., 2020). These disruptions in neural function, specifically slowing of neural transmission, may exert effects on cognitive processing speed (Zhao et al., 2021).

There is also compelling evidence for associations between glial structures that mediate glymphatic function via astrocyte AQP4 channels and cognition in both healthy (Hutchison et al., 2013; Abdelkarim et al., 2019; Zeppenfeld et al., 2017) and pathological aging (Sivakolundu et al., 2020; Wang et al., 2022; Zeppenfeld et al., 2017). Astrocyte AQP4 channels are necessary for exchange between brain interstitial fluid and CSF in the perivascular space (Iliff et al., 2012). They also mediate neurovascular coupling processes that are adversely affected in aging (Abdelkarim et al., 2019; West et al., 2019; Turner et al., 2022; Zimmerman et al., 2021; Tarantini et al., 2017) and may lead to reduced processing speed (Zhao et al., 2021). Conversely, it is also possible that impaired glymphatic clearance may impact neurovascular coupling, resulting in processing speed variations among healthy individuals (Abdelkarim et al., 2019; Zhao et al., 2021; Zimmerman et al., 2021). Such processes could then exert broader deleterious influences on higher-order cognition (Salthouse, 1996; Schubert et al., 2023; Motes et al., 2011) and should be explored in future studies.

While our study has limitations, we nonetheless demonstrated significant relationships between perivascular CSF flow, meningeal lymphatic vessels, and processing speed. These results support the hypothesis that age-related declines in glymphatic integrity play an important role in cognitive aging, specifically, processing speed declines. More work with larger sample sizes and methods that are considered more direct measurements of glymphatic function, including CSF fraction (Zhou et al., 2024), 3D-T1 spatial sequences designed for mLV visualization (Zhou et al., 2024), and gadolinium-based contrast agents (GBCA) via intrathecal administration (Zhang et al., 2021; Wu et al., 2021), should be employed in future studies to further test this hypothesis.

Overall, this study suggests intriguing connections between CSF flow, waste clearance structures in the CNS, and their impact on brain function, particularly processing speed. We found that the efficiency of glymphatic function declines with age, which may lead to compensatory increases in the volume of the efflux structure (mLVs) of the system and variations in processing speed in healthy individuals. Our findings regarding the glymphatic system and mLVs have implications for the neurophysiologic basis of age-related differences in cognition.

## Acknowledgements

We thank all the patients and volunteers who participated in this research efforts. We thank our lab research interns and coordinators for their help in recruiting patients and volunteering for the study. This work was supported by a National Multiple Sclerosis Society Research Grant (RG-1507-04951) to BR.

## References

Abdelkarim D, Zhao Y, Turner MP, Sivakolundu DK, Lu H, Rypma B (2019) A neural-vascular complex of age-related changes in the human brain: Anatomy, physiology, and implications for neurocognitive aging. Neuroscience & Biobehavioral Reviews.

Absinta M, Ha S-K, Nair G, Sati P, Luciano NJ, Palisoc M, Louveau A, Zaghloul KA, Pittaluga S, Kipnis J, Reich DS (2017) Human and nonhuman primate meninges harbor lymphatic vessels that can be visualized noninvasively by MRI Johansen-Berg H, ed. eLife 6:e29738.

Ahn JH, Cho H, Kim JH, Kim SH, Ham JS, Park I, Suh SH, Hong SP, Song JH, Hong YK, Jeong Y, Park SH, Koh GY (2019) Meningeal lymphatic vessels at the skull base drain cerebrospinal fluid. Nature. 572:62–66.

Albayram MS, Smith G, Tufan F, Tuna IS, Bostancıklıoğlu M, Zile M, Albayram O (2022) Non-invasive MR imaging of human brain lymphatic networks with connections to cervical lymph nodes. Nat Commun 13:203.

Alshammari, Fawaz & Keil, Samantha & Wilcox, Mary. (2026). Sleep fragmentation, impaired glymphatic clearance, and long-term cognitive impairment after critical illness. Critical Care Science. 38. 10.62675/2965-2774.20260042.

Andersson JLR, Graham MS, Zsoldos E, Sotiropoulos SN (2016) Incorporating outlier detection and replacement into a non-parametric framework for movement and distortion correction of diffusion MR images. NeuroImage 141:556–572.

Andersson JLR, Sotiropoulos SN (2016) An integrated approach to correction for off-resonance effects and subject movement in diffusion MR imaging. NeuroImage 125:1063–1078.

Antila S, Karaman S, Nurmi H, Airavaara M, Voutilainen MH, Mathivet T, Chilov D, Li Z, Koppinen T, Park J-H, Fang S, Aspelund A, Saarma M, Eichmann A, Thomas J-L, Alitalo K (2017) Development and plasticity of meningeal lymphatic vessels. Journal of Experimental Medicine 214:3645–3667.

Aspelund A, Antila S, Proulx ST, Karlsen TV, Karaman S, Detmar M, Wiig H, Alitalo K (2015) A dural lymphatic vascular system that drains brain interstitial fluid and macromolecules. Journal of Experimental Medicine 212:991–999.

Avants BB, Tustison NJ, Song G, Cook PA, Klein A, Gee JC (2011) A reproducible evaluation of ANTs similarity metric performance in brain image registration. Neuroimage 54:2033–2044.

Carare, R. O., Aldea, R., Agarwal, N., Bacskai, B. J., Bechman, I., Boche, D., Bu, G., Bulters, D., Clemens, A., Counts, S. E., de Leon, M., Eide, P. K., Fossati, S., Greenberg, S. M., Hamel, E., Hawkes, C. A., Koronyo-Hamaoui, M., Hainsworth, A. H., Holtzman, D., … Verma, A. (2020). Clearance of interstitial fluid (ISF) and CSF (CLIC) group-part of Vascular Professional Interest Area (PIA): Cerebrovascular disease and the failure of elimination of Amyloid-β from the brain and retina with age and Alzheimer’s disease-Opportunities for Therapy. Alzheimer’s & Dementia, 12(1), e12053. 10.1002/dad2.12053.

Carotenuto A, Cacciaguerra L, Pagani E, Preziosa P, Filippi M, Rocca MA (2022) Glymphatic system impairment in multiple sclerosis: relation with brain damage and disability. Brain 145:2785–2795.

Cheung, G. W., & Lau, R. S. (2008). Testing mediation and suppression effects of latent variables: Bootstrapping with structural equation models. Organizational Research Methods, 11(2), 296–325. 10.1177/1094428107300343

Da Mesquita S, Louveau A, Vaccari A, Smirnov I, Cornelison RC, Kingsmore KM, Contarino C, Onengut-Gumuscu S, Farber E, Raper D, Viar KE, Powell RD, Baker W, Dabhi N, Bai R, Cao R, Hu S, Rich SS, Munson JM, Lopes MB, Overall CC, Acton ST, Kipnis J (2018) Functional aspects of meningeal lymphatics in ageing and Alzheimer’s disease. Nature 560:185–191.

Dagum, P., Elbert, D. L., Giovangrandi, L., Singh, T., Venkatesh, V. V., Corbellini, A., Kaplan, R. M., Levendovszky, S. R., Ludington, E., Yarasheski, K., Lowenkron, J., VandeWeerd, C., Lim, M. M., & Iliff, J. J. (2026). The glymphatic system clears amyloid beta and tau from brain to plasma in humans. Nature communications, 17(1), 715. 10.1038/s41467-026-68374-8.

Ding X-B, Wang X-X, Xia D-H, Liu H, Tian H-Y, Fu Y, Chen Y-K, Qin C, Wang J-Q, Xiang Z, Zhang Z-X, Cao Q-C, Wang W, Li J-Y, Wu E, Tang B-S, Ma M-M, Teng J-F, Wang X-J (2021) Impaired meningeal lymphatic drainage in patients with idiopathic Parkinson’s disease. Nat Med 27:411–418.

Eckstein K, Bachrata B, Hangel G, Widhalm G, Enzinger C, Barth M, Trattnig S, Robinson SD (2021) Improved susceptibility weighted imaging at ultra-high field using bipolar multi-echo acquisition and optimized image processing: CLEAR-SWI. NeuroImage 237:118175.

Hein, Z. M., Che Mohd Nassir, C. M. N., Abdul Hamid, H., Abdullah, M. F. I. L., & Shantakumari, N. (2026). The Prevalence of Sleep Disorders in Populations with Glymphatic Dysfunction: A Systematic Review and Meta-Analysis. Biology, 15(4), 309. 10.3390/biology15040309.

Hutchison JL, Lu H, Rypma B (2013) Neural mechanisms of age-related slowing: the ΔCBF/ΔCMRO2 ratio mediates age-differences in BOLD signal and human performance. Cereb Cortex 23(10):2337–46.

Iliff JJ, Wang M, Liao Y, Plogg BA, Peng W, Gundersen GA, Benveniste H, Vates GE, Deane R, Goldman SA, Nagelhus EA, Nedergaard M (2012) A Paravascular Pathway Facilitates CSF Flow Through the Brain Parenchyma and the Clearance of Interstitial Solutes, Including Amyloid β. Science Translational Medicine 4:147ra111–147ra111.

Jessen NA, Munk ASF, Lundgaard I, Nedergaard M (2015) The Glymphatic System: A Beginner’s Guide. Neurochem Res 40:2583–2599.

Jiang Q, Zhang L, Ding G, Davoodi-Bojd E, Li Q, Li L, Sadry N, Nedergaard M, Chopp M, Zhang Z (2017) Impairment of the glymphatic system after diabetes. J Cereb Blood Flow Metab 37:1326–1337.

Kress BT, Iliff JJ, Xia M, Wang M, Wei HS, Zeppenfeld D, Xie L, Kang H, Xu Q, Liew JA, Plog BA, Ding F, Deane R, Nedergaard M (2014) Impairment of paravascular clearance pathways in the aging brain. Annals of Neurology 76:845–861.

Lee H-J, Lee DA, Shin KJ, Park KM (2022) Glymphatic system dysfunction in obstructive sleep apnea evidenced by DTI-ALPS. Sleep Medicine 89:176–181.

Li L, Ding G, Zhang L, Davoodi-Bojd E, Chopp M, Li Q, Zhang ZG, Jiang Q (2022) Aging-Related Alterations of Glymphatic Transport in Rat: In vivo Magnetic Resonance Imaging and Kinetic Study. Front Aging Neurosci 14:841798.

Louveau A, Smirnov I, Keyes TJ, Eccles JD, Rouhani SJ, Peske JD, Derecki NC, Castle D, Mandell JW, Lee KS, Harris TH, Kipnis J (2015) Structural and functional features of central nervous system lymphatic vessels. Nature 523:337–341.

Mestre H et al. (2020) Cerebrospinal fluid influx drives acute ischemic tissue swelling. Science 367:eaax7171.

Motes MA, Biswal BB, Rypma B (2011) Age-Dependent Relationships between Prefrontal Cortex Activation and Processing Efficiency. Cogn Neurosci 2(1):1–10.

Nedergaard, M., & Goldman, S. A. (2020). Glymphatic failure as a final common pathway to dementia. *Science (New York*, N.Y.), 370(6512), 50–56. 10.1126/science.abb8739.

R Core Team. (2018). R: A language and environment for statistical computing (Version 3.5.2) [Computer software]. R Foundation for Statistical Computing. https://www.R-project.org/.

Rosseel Y (2012) lavaan: An R Package for Structural Equation Modeling. Journal of Statistical Software 48:1–36.

Rypma B, Berger JS, Prabhakaran V, Martin Bly B, Kimberg DY, Biswal BB, D’Esposito M (2006) Neural correlates of cognitive efficiency. NeuroImage 33:969–979.

Salthouse TA (1996) The processing-speed theory of adult age differences in cognition. Psychological Review 103:403–428.

Salthouse, T. A. (2000). Aging and measures of processing speed. Biological Psychology, 54(1), 35–54. 10.1016/S0301-0511(00)00052-1.

Schubert AL, Löffler C, Hagemann D, Sadus K (2023) How robust is the relationship between neural processing speed and cognitive abilities? Psychophysiology 60(2):e14165.

Sivakolundu DK, West KL, Maruthy GB, Zuppichini M, Turner MP, Abdelkarim D, Zhao Y, Nguyen D, Spence JS, Lu H, Okuda DT, Rypma B (2019) Reduced arterial compliance along the cerebrovascular tree predicts cognitive slowing in multiple sclerosis: Evidence for a neurovascular uncoupling hypothesis. Multiple Sclerosis Journal:135245851986660.

Sivakolundu DK, West KL, Zuppichini M, Turner MP, Abdelkarim D, Zhao Y, Spence JS, Lu H, Okuda DT, Rypma B (2020) The neurovascular basis of processing speed differences in humans: A model-systems approach using multiple sclerosis. NeuroImage:116812.

Taoka T, Masutani Y, Kawai H, Nakane T, Matsuoka K, Yasuno F, Kishimoto T, Naganawa S (2017) Evaluation of glymphatic system activity with the diffusion MR technique: diffusion tensor image analysis along the perivascular space (DTI-ALPS) in Alzheimer’s disease cases. Jpn J Radiol 35:172–178.

Tarantini S, Tran CHT, Gordon GR, Ungvari Z, Csiszar A (2017) Impaired neurovascular coupling in aging and Alzheimer’s disease: Contribution of astrocyte dysfunction and endothelial impairment to cognitive decline. Exp Gerontol 94:52–58.

Tatekawa H, Matsushita S, Ueda D, Takita H, Horiuchi D, Atsukawa N, Morishita Y, Tsukamoto T, Shimono T, Miki Y (2023) Improved reproducibility of diffusion tensor image analysis along the perivascular space (DTI-ALPS) index: an analysis of reorientation technique of the OASIS-3 dataset. Jpn J Radiol 41:393–400.

Turner MP, Zhao Y, Abdelkarim D, Liu P, Spence JS, Hutchison JL, Sivakolundu DK, Thomas BP, Hubbard NA, Xu C, Taneja K, Lu H, Rypma B (2022) Altered linear coupling between stimulus-evoked blood flow and oxygen metabolism in the aging human brain. Cereb Cortex 33:135–151.

Verheggen, I. C. M., Van Boxtel, M. P. J., Verhey, F. R. J., Jansen, J. F. A., & Backes, W. H. (2018). Interaction between blood-brain barrier and glymphatic system in solute clearance. Neuroscience and biobehavioral reviews, 90, 26–33. 10.1016/j.neubiorev.2018.03.028

Wang J, Zhou Y, Zhang K, Ran W, Zhu X, Zhong W, Chen Y, Li J, Sun J, & Lou M. (2023) Glymphatic function plays a protective role in ageing-related cognitive decline. Age and Ageing 52(7):afad107.

Wang L, Zhang Y, Zhao Y, Marshall C, Wu T, Xiao M (2019) Deep cervical lymph node ligation aggravates AD-like pathology of APP/PS1 mice. Brain Pathol 29(2):176–92.

Wang Y, Huang C, Guo Q, Chu H (2022) Aquaporin-4 and Cognitive Disorders. Aging Dis. 13(1):61–72.

West KL, Zuppichini MD, Turner MP, Sivakolundu DK, Zhao Y, Abdelkarim D, Spence JS, Rypma B (2019) BOLD hemodynamic response function changes significantly with healthy aging. NeuroImage 188:198–207.

Wu, CH, Lirng, JF, Ling, YH, Wang, YF, Wu, HM, Fuh, JL, Lin, PC, Wang, SJ, & Chen, SP (2021) Noninvasive Characterization of Human Glymphatics and Meningeal Lymphatics in an in vivo Model of Blood-Brain Barrier Leakage. Annals of neurology. 89(1):111–12.

Xiong Y, Yu Q, Zhi H, Peng H, Xie M, Li R, Li K, Ma Y, Sun P (2024) Advances in the study of the glymphatic system and aging. CNS Neurosci Ther. 30(6):e14803.

Zeppenfeld DM, Simon M, Haswell JD, D’Abreo D, Murchison C, Quinn JF, Grafe MR, Woltjer RL, Kaye J, Iliff JJ (2017) Association of Perivascular Localization of Aquaporin-4 With Cognition and Alzheimer Disease in Aging Brains. JAMA Neurol. 74(1):91–99.

Zhang W, Zhou Y, Wang J, Gong X, Chen Z, Zhang X, Cai J, Chen S, Fang L, Sun J, Lou M (2021) Glymphatic clearance function in patients with cerebral small vessel disease. NeuroImage 238:118257.

Zhao Y, Liu P, Turner, MP, Abdelkarim, D, Lu, H, & Rypma, B (2021) The neural-vascular basis of age-related processing speed decline. Psychophysiology, 58(7), e13845.

Zhou L, Nguyen TD, Chiang GC, Wang XH, Xi K, Hu TW, Tanzi EB, Butler TA, de Leon MJ, Li Y (2024) Parenchymal CSF fraction is a measure of brain glymphatic clearance and positively associated with amyloid beta deposition on PET Alzheimers Dement. Alzheimer’s & dementia. 20(3):2047–2057.

Zhou W, Shen B, Shen W, Chen H, Zheng Y and Fei J (2020) Dysfunction of the Glymphatic System Might Be Related to Iron Deposition in the Normal Aging Brain. Front. Aging Neurosci. 12:559603. doi: 10.3389/fnagi.2020.559603.

Zimmerman B, Rypma B, Gratton G, Fabiani M (2021) Age-related changes in cerebrovascular health and their effects on neural function and cognition: A comprehensive review. Psychophysiology 58(7):e13796.

